# Genomic signatures associated with the evolutionary loss of egg yolk in parasitoid wasps

**DOI:** 10.1101/2023.12.30.573744

**Authors:** Xianxin Zhao, Yuanyuan Liu, Yi Yang, Chun He, Kevin C. Chan, Haiwei Lin, Qi Fang, Gongyin Ye, Xinhai Ye

## Abstract

Trait regression and loss have occurred repeatedly in numerous lineages throughout evolutionary history in response to changing environments. In parasitoid wasps, a mega-diverse group of hymenopteran insects, loss or reduction of yolk in the egg has been reported in many species. This phenotypic change likely evolved as a response to the shift from ectoparasitism to endoparasitism. However, the genetic basis of this trait and the impact of its loss on genome evolution remain poorly understood. Here, we performed a comparative genomic analysis of 64 hymenopteran insects. The conserved insect yolk protein gene *vitellogenin* (*Vg*) underwent five independent loss events in four families, involving 23 of the analyzed endoparasitoid species. Whole-genome alignment suggested that *Vg* loss occurred during genome rearrangement events. Analysis of *Vg* receptor gene (*VgR*) loss, selection, and structural variation in lineages lacking *Vg* demonstrated functional biases in the patterns of gene loss. The ectoparasitism to endoparasitism transition did not appear to be the primary driver of *Vg* loss or the subsequent *VgR* evolution. A number of parallel and convergent genomic changes were observed between *Vg*-loss lineages, including gene family evolution and selection of genes related to transport, development, and metabolism. These changes may have facilitated embryonic development without the yolk in these lineages. Together, these findings reveal the genomic basis underlying a unique trait loss in parasitoid wasps. More broadly, this study enhances our understanding of yolk loss evolution outside the class Mammalia, highlighting a potential evolutionary trend arising from the availability of an alternative nutrient source for embryonic development.

## Introduction

A fundamental goal in the field of evolutionary biology is to understand the genetic bases underlying phenotypic shifts and evolutionary novelties throughout the tree of life. Trait regression or loss (the reduction or complete loss, respectively, of an ancestral trait) are phenotypic shifts that have occurred repeatedly throughout evolutionary history. These shifts likely occur in response to alterations in the environment, habitat, or species interaction network^1–3^. Multiple genome sequencing and comparative genomic studies have been conducted to uncover the genetic bases of evolutionary trait loss in animals, including eye and pigmentation loss among animals living in darkness^4–8^, tooth loss in birds^9–11^, limb loss in snakes^12^, and tail loss in humans and apes^13^. Such studies have highlighted the importance of gene loss and inactivating mutations in the evolution of trait loss or regression. These investigations have also demonstrated the power of comparative genomic approaches to reveal the genetic mechanisms underlying complex trait evolution.

Trait loss and the associated genetic bases are topics of great research interest in the context of parasite evolution, particularly because long-term host dependence may lead to trait loss in processes such as nutrient acquisition^14,15^. For example, many parasitic plants show a loss of genes associated with photosynthesis, nutrient uptake, flowering regulation, defense responses, and root development^16–19^. A large-scale genomic analysis of parasitic nematodes and platyhelminths revealed the widespread losses of metabolic pathways and differential pathway coverage across different evolutionary clades^20^. Other studies have revealed the loss of homeobox gene families^21^ and cilium-related genes^22^ in parasitic worms, which may be related to their simplified morphology.

Insects are the largest and most diverse group of organisms on Earth. Parasitoids, defined as parasitic species that eventually kill their hosts, have evolved repeatedly and independently in several insect orders^23^. The most common and representative parasitoid species are parasitoid wasps of the order Hymenoptera; these are extremely biologically diverse, and a rapid accumulation of genomic data has made them model organisms in the field of evolutionary biology^24–31^. Comparative genomic analyses of parasitoid wasps and their free-living relatives can reveal novel evolutionary trait losses or regressions, providing new insights into the evolution of parasitism.

Some parasitoid wasp species exhibit an intriguing trait loss with a visible phenotype: loss of the egg yolk. Eggs with little or no yolk were first recorded in the endoparasitoid *Chalcis sispes*, and are widespread among parasitoid wasps from various lineages^33–37^ (Supplementary Table 1). Such yolkless eggs have been termed “hydropic”, referring to their method of nutrient acquisition from host hemolymph, which distinguishes them from anhydropic eggs^32^. Moreover, the phenomenon of radiolabeled amino acids absorption from host hemolymph was found in *Microplitis croceipes*, rather than protein taken^33^, implying that yolkless insect eggs have evolved to obtain the nutrients necessary for development directly from the host. Indeed, yolk trait loss has been likely linked to the evolution of parasitic strategies (e.g., the shift from ectoparasitism to endoparasitism). Thus, yolk loss has likely evolved as a result of embryonic development inside the host^23,38,39^, although this has not been proven.

To uncover the genetic basis of egg yolk loss in parasitoid wasps, we here conducted a comparative genomic analysis using *vitellogenin* (*Vg*), a conserved gene encoding an egg yolk protein precursor in insects^38,40^. Using a broad, representative selection of hymenopteran species, we identified lineages with *Vg* loss events and established a putative loss mechanism. Furthermore, we examined loss of a *Vg*-related gene to assess co-evolution in multiple lineages. These findings enhance our knowledge of yolk loss and parasitoid wasp genome evolution, contributing more broadly to an improved understanding of the genomic changes associated with shifts in parasitic strategies and embryonic developmental conditions.

## Results

### *Vg* loss was recurrent in Hymenoptera but confined to endoparasitoid wasps

We first sought to identify *Vg*s and homologous genes in members of the order Hymenoptera. Thus, a comparative genomic investigation was conducted in 64 representative hymenopteran species, including the major lineages: sawflies, wasps, ants, and bees (Fig. 1b). A comprehensive homology-based search was followed by protein domain screening and manual curation (see **Methods**), yielding a total of 237 genes. These genes were categorized into three groups based on the presence or absence of specific functional domains: 64 complete classic *Vg* genes (*Vg*s), 169 *Partial Vg* genes (*PVg*s), and four *Vg-like* genes (*Vgl*s) (Fig. 1a; Supplementary Table 2). The classic *Vg*s encoded proteins with three major domains^38,41^, namely Vitellogenin_N or LPD_N (PF01347), Vit_openb-sht (PF09172), and von Willebrand factor type D (vWD) (PF00094); notably, we identified seven *Vg*s in seven species that were incorrectly annotated in prior genome annotations (Supplementary Fig. 1). The *PVg*s encoded other proteins containing the Vitellogenin_N domain, whereas *Vgl*s encoded proteins without the Vitellogenin_N domain but with the Vit_openb-sht and/or vWD domain.

**Fig. 1.**
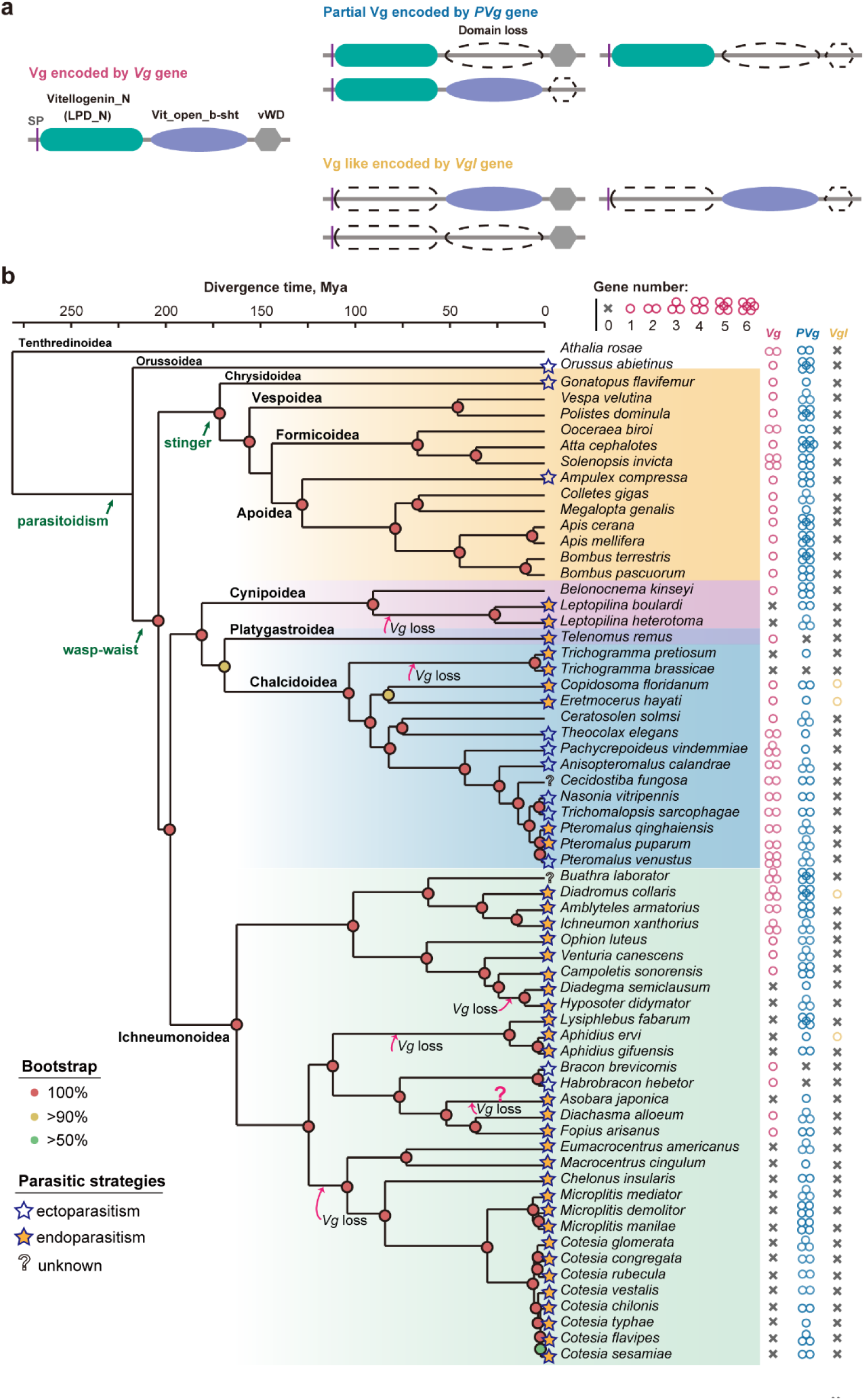
Evolutionary analyses of vitellogenin (*Vg*) and *Vg*-related genes. (**a**) Domain schematics of Vg and homologs, namely Partial Vg (PVg) and Vg-like (Vgl) proteins. Vg proteins had three major domains: Vitellogenin_N (LPD_N) (green), Vit_open_b-sht (purple), and von Willebrand factor type D (vWD) (gray). Other Vg-related proteins were classified as PVgs or Vgls based on the presence or absence, respectively, of the Vitellogenin_N domain. SP, signal peptide. (**b**) Phylogenic analysis of hymenopteran insects. The tree suggested the existence of at least five independent *Vg* loss events. Divergence times were estimated using 12 calibration points. The colored solid circle on each tree node indicates the bootstrap value. Stars to the left of the species name indicate the parasitic strategy of that species, endoparasitism (yellow) or ectoparasitism (white). Question marks to the left of the species name indicate an unknown parasitic strategy. The numbers of *Vg* and *Vg*- related genes in each genome are indicated at right; the pink question mark for *Vg* in *Asobara japonica* indicates ambiguity. This loss may have been an artifact from genome assembly (Supplementary Fig. 2).

To place our findings in an evolutionary context, a phylogenetic tree was constructed for the 64 species using 1116 single-copy orthologous genes. The *Vg* identification results were then mapped onto the resulting tree. This revealed a great deal of variation in *Vg* copy number within the order Hymenoptera. Interestingly, *Vg* was absent from the genomes of 24 parasitoid wasps in three superfamilies (Ichneumonoidea, Chalcidoidea and Cynipoidea) that contain diverse parasitoid species (Fig. 1b; Supplementary Table 2). In contrast, one to four copies of *Vg* were conserved in all other non-parasitoid hymenopteran species, including sawfly (*Athalia*), vespid wasps (*Vespa* and *Polistes*), ants (*Ooceraea*, *Atta*, and *Solenopsis*), and bees (*Colletes*, *Megalopta*, *Apis*, and *Bombus*). *Vg* loss was presumed in the *Drosophila* parasitoid *Asobara japonica* (Braconidae), but this may have been an artifact due to the presence of a *PVg* gene located at the end of the scaffold (Supplementary Fig. 2; Fig. 1b).

*Vg* loss appeared to have occurred independently in five different parasitoid wasp lineages, leading to the observed lack of *Vg* in 23 species (excluding *A. japonica*). Specifically, *Vg* was lost in the common ancestor of two *Leptopilina* species in the superfamily Cynipoidea; the common ancestor of two *Trichogramma* species in the superfamily Chalcidoidea; the common ancestor of *Diadegma semiclausum* and *Hyposoter didymator* in the family Ichneumonidae; the common ancestor of three aphid parasitoids of the genera *Aphidius* and *Lysiphlebus* in the family Braconidae; and the common ancestor of 14 braconid parasitoids from the genera *Cotesia*, *Microplitis*, *Chelonus*, *Macrocentrus*, and *Eumacrocentrus* in the family Braconidae (Fig. 1b). In addition, the phylogenetic tree demonstrated extensive variations in *PVg* and *Vgl* gene copy number across hymenopteran species, suggesting complex evolutionary histories.

To understand the relationships between *Vg* loss and parasitic strategies, the 64 species were divided into three categories based on the location in which their eggs are laid and embryos develop: ectoparasitoids (those that lay eggs on the external host surface), endoparasitoids (those that lay eggs inside the host)^23^, and unknown.

These categorizations were then mapped to the phylogenic tree (Fig. 1b). Based on the key role of Vg proteins in yolk formation (vitellogenesis)^40^, we hypothesized that *Vg* losses would be restricted to endoparasitoids due to the nutrient-rich environment (host hemolymph) ^42,43^ in which the eggs reside. Indeed, all of the parasitoid wasps with *Vg* loss were endoparasitoids. However, not all of the analyzed endoparasitoids demonstrated *Vg* loss; 35% retained copies of *Vg*. This suggested that endoparasitism may not have been the primary factor driving *Vg* loss in parasitoid wasps.

### *PVg* was not derived from *Vg* disruption in *Vg*-loss species

We hypothesized that recently mutated *Vg*s retaining the Vitellogenin_N domain (i.e., *PVg*s) may have retained some *Vg* functionality due to the importance of the N-domain for oocyte uptake^44,45^. Thus, *Vg* loss may not have directly resulted in a lack of vitellogenesis if *PVg* was present. To investigate the evolutionary origins of *PVg*s and to test whether these genes arose from recent *Vg* disruption among *Vg*-loss species, a phylogenetic analysis was performed using all *Vg*s and *PVg*s identified in this study. If the *PVg*s had primarily originated from *Vg* mutation, *PVg*s would cluster together with *Vg*s. Instead, the Vg sequences showed a distinct evolutionary history from the PVg sequences (Fig. 2a). Notably, the N-terminal domain sequences of eight PVgs in seven species were located in the Vg clade, suggesting that these specific PVgs may have resulted from Vg disruption. However, none of the corresponding species had undergone *Vg* loss events, with the exception of the ambiguous *Vg* loss in *A. japonica* as noted above.

**Fig. 2.**
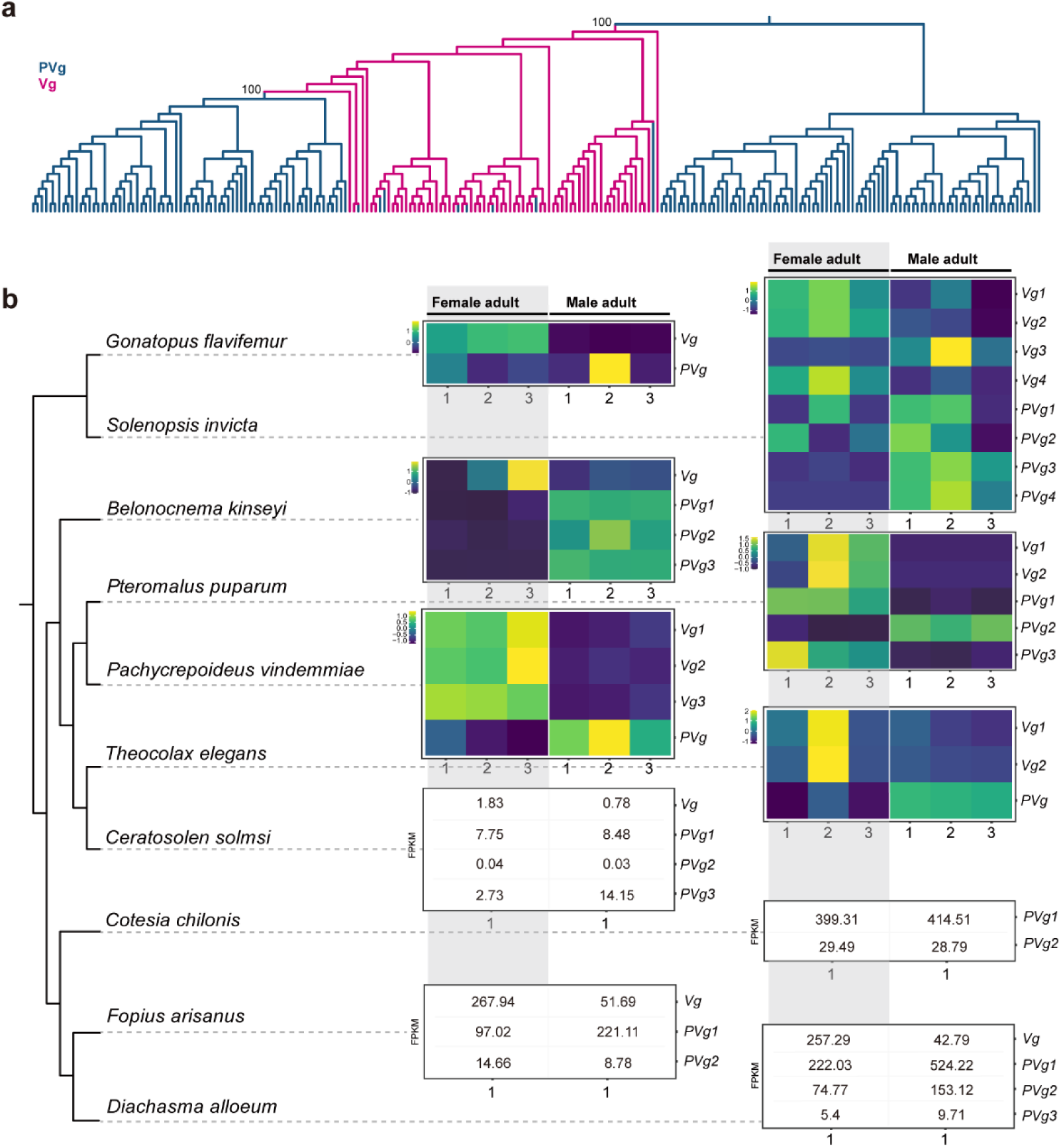
Phylogenetic and gene expression analyses of *Vg*s and *PVg*s. (**a**) Phylogenetic tree of *Vg*s (red) and *PVg*s (blue) in hymenopteran species. All of the *Vg*s clustered together with some of the *PVg*s (∼48%); the remainder of the *PVg*s clustered together as a monophyletic group. (**b**) Expression patterns of *Vg*s and *PVg*s in male and female adults of 10 hymenopteran species. Columns indicate individual samples; each row represents expression of a *Vg* or *PVg* (as noted at right). Expression values are represented with color and were calculated in fragments per kilobase of transcript per million mapped reads (FPKM). For experiments without biological replicates, gene expression values are shown in numerical form. The phylogeny of the 10 selected hymenopteran insects was extracted from the data shown in Fig. 1.

The functions of *PVg*s in insects have not yet been characterized. To establish whether there was putative functional overlap between *Vg*s and *PVg*s, we examined *Vg* and *PVg* expression in 10 hymenopteran species for which there were publicly available RNA sequencing data generated from adults of both sexes. *Vg* was generally expressed at higher levels in females than in males, whereas the opposite was true of *PVg* (Fig. 2b; Supplementary Table 3). These distinct expression patterns between *Vg*s and *PVg*s in adult hymenopterans suggested that the genes have divergent functionality. Overall, these results revealed dramatic differences between *Vg*s and *PVg*s with respect to both evolutionary history and gene expression. This supported the hypothesis that vitellogenesis was largely arrested in endoparasitoid wasps with *Vg* loss, consistent with previous reports of yolkless eggs^32,35^ (Supplementary Table 1).

### *Vg* loss was likely caused by genome rearrangement

To explore the potential causes of *Vg* loss in endoparasitoid wasps, the genomes of species with *Vg* loss were examined for regions that were syntenic with *Vg* loci in their *Vg*-retaining relatives. Because *Vg* loss events span large evolutionary distances across hymenopteran phylogeny, syntenic regions containing *Vg* were examined only within each family or superfamily. In Ichneumonidae, the genes encoding chondroitin sulfate N-acetylgalactosaminyltransferase and alanine-glyoxylate aminotransferase were identified as marker genes for the *Vg* region; these two genes flanked the genomic region containing *Vg* in wasps of the *Vg*- retaining genera *Ophion* and *Venturia*. A search for remnants of *Vg* (e.g., a pseudogene or inactive gene) in the corresponding region of *H. didymator*, which had lost *Vg*, revealed no homologous sequences (Fig. 3a; see **Methods**). Similar searches did not reveal *Vg* remnants in other *Vg*-loss species of the families/superfamilies Braconidae, Cynipoidea, or Chalcidoidea (Fig. 3b, c; Supplementary Fig. 3). These results suggested that *Vg* loss was not due to gene-inactivating mutations. However, it is also possible that rapid sequence evolution prevented homologous sequence detection.

**Fig. 3.**
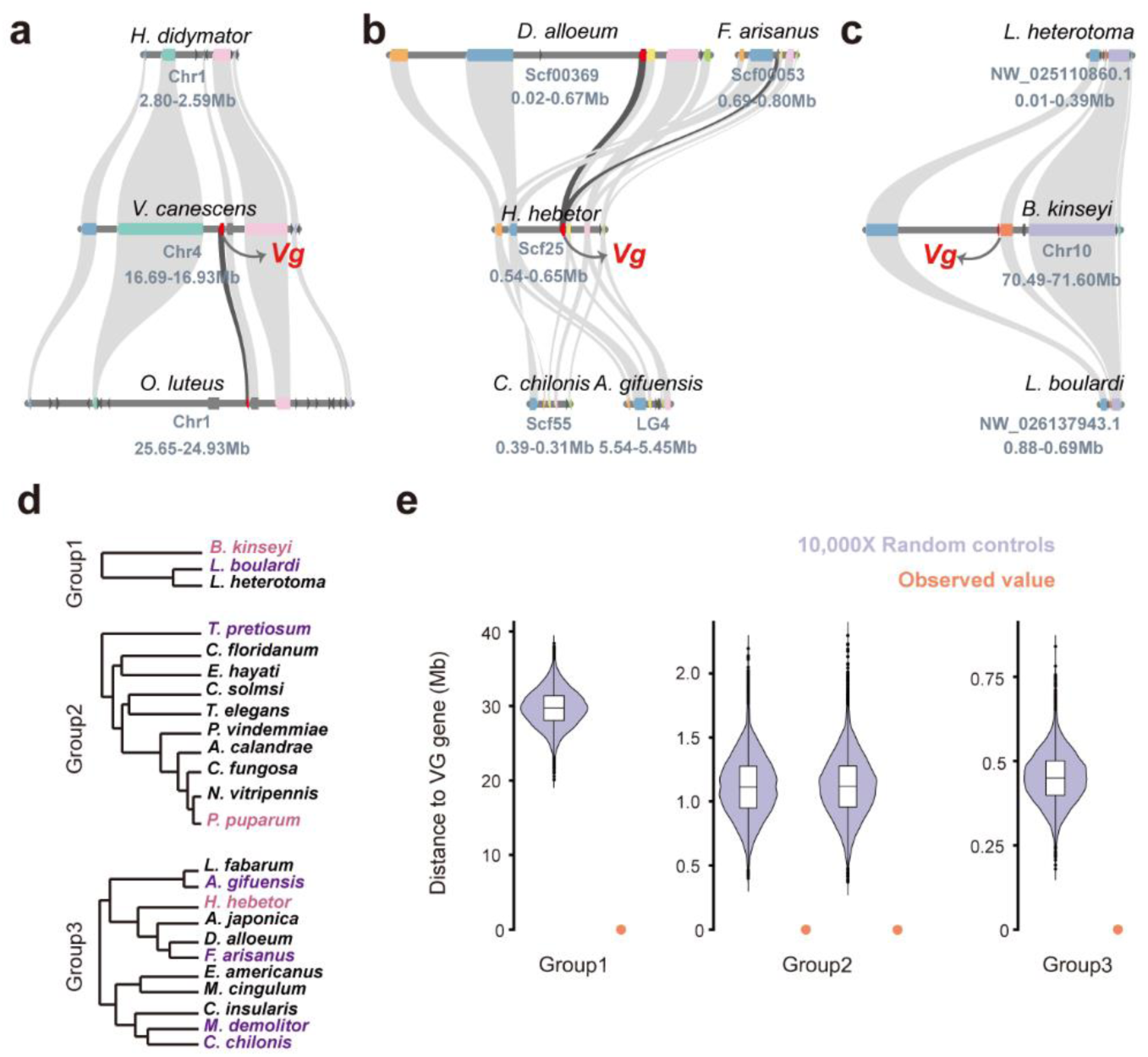
*Vg* was likely lost from endoparasitoid Hymenoptera due to genome rearrangement. (**a–c**) Synteny of *Vg* and the flanking genes among representative *Vg*-loss and *Vg*-retaining species within the (**a**) family Ichneumonidae, (**b**) family Braconidae, and (**c**) superfamily Cynipoidea. *Vg*s are indicated with red. Grey lines indicate orthologous relationships. (**d**) Phylogenetic analyses of three independent lineages with *Vg* loss. Group1, Cynipoidea; Group2, Chalcidoidea; Group3, Braconidae. Pairwise genome alignments were performed within each group to identify genomic rearrangement breakpoints. Species shown in pink and purple were *Vg*-retaining (reference) and *Vg*-loss (query) species, respectively. (**e**) Distance between *Vg* loci and observed genomic rearrangement breakpoints (orange points) or randomly shuffled control bins (purple violins).

We next asked whether genome rearrangements had contributed to *Vg* losses. This was addressed through detection of genome rearrangement breakpoints between *Vg*-loss species and their *Vg*-retaining relatives in three independent lineages (Cynipoidea, Chalcidoidea, and Braconidae) via whole-genome alignment (Fig. 3d; see **Methods**). Ichneumonid species were excluded from this analysis because the genome assemblies were highly fragmented, with over 13,000 scaffolds for *Vg*-loss species, which could have negatively affected the accuracy of genome-wide alignments. In Cynipoidea (Group 1), 2151 genome rearrangement breakpoints were identified between *Leptopilina boulardi* (*Vg* loss) and *Belonocnema kinseyi* (*Vg* retaining). Notably, the breakpoints significantly overlapped with the *Vg* locus in the *B. kinseyi* genome (Fig. 4e). Similar results were observed for species in Chalcidoidea and Braconidae (Fig. 4e; Observed breakpoints vs. a set of control regions by 10,000 times randomly shuffling breakpoints on the same chromosome, *p* = 0, one-tailed permutation test). Taken together, these findings indicated strong contributions of genome rearrangement to *Vg* loss events in endoparasitoid wasps.

**Fig. 4.**
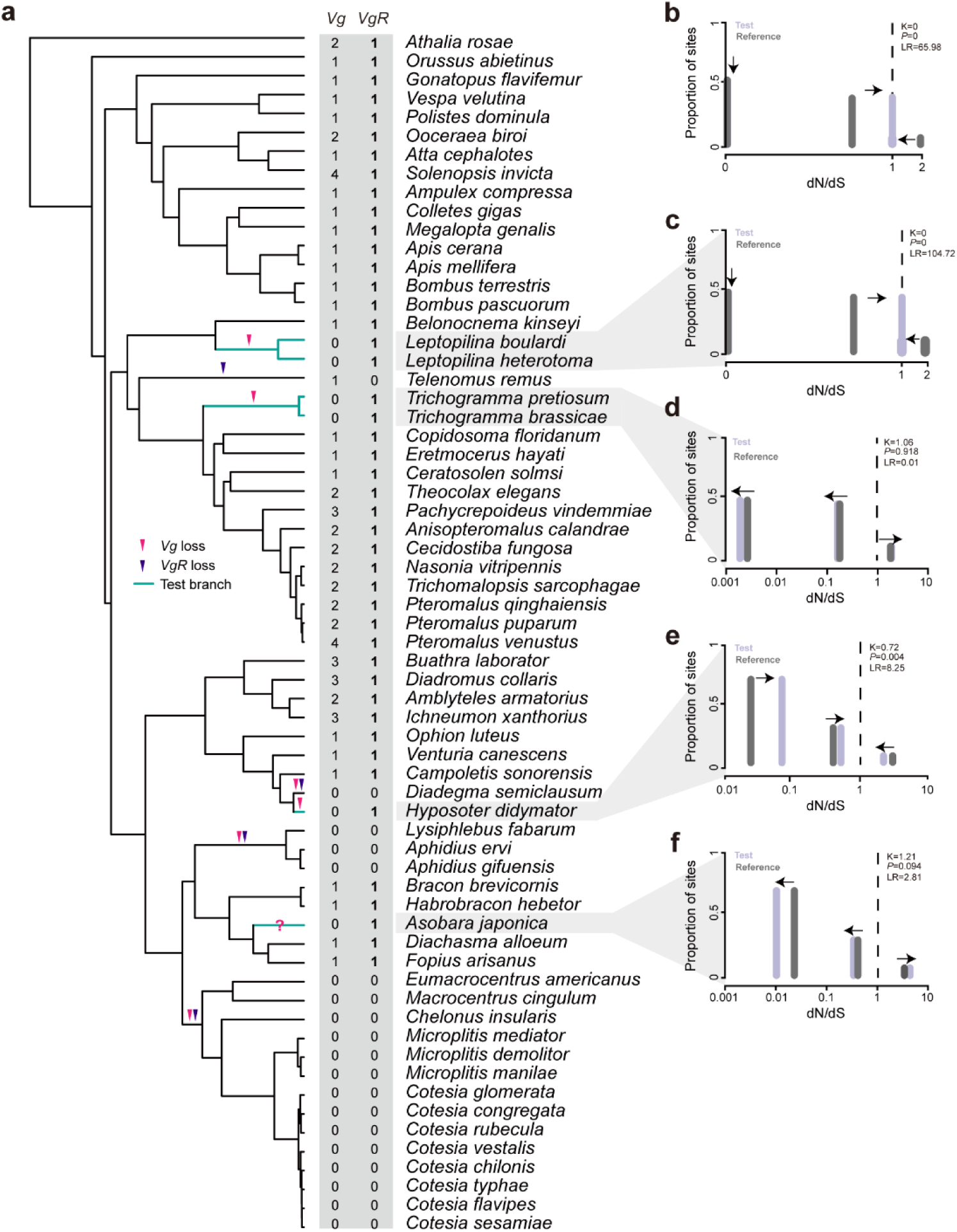
Relaxed selection and complete loss of genes encoding the Vg receptor (*VgR*) in many lineages with *Vg* loss. (**a**) Phylogenetic tree (from Fig. 1b) showing instances of *Vg* and *VgR* loss (pink and purple arrows, respectively). Branches shown in green were used in molecular selection analyses. (**b–f**) Selection signatures of *VgR*s from independent analyses of several *Vg*-loss branches or clades, namely (**b**) all *Vg*- loss branches or clades, excluding *Asobara japonica*; (**c**) *Leptopilina* spp.; (**d**) *Trichogramma* spp.; (**e**) *Hyposoter didymator*; and (**f**) *A. japonica*. Nonsynonymous to synonymous substitution rate (dN/dS) distributions across sites in the *VgR*s are shown for test (branches listed above in **b–f**, light purple) and reference (branches left from test branches, dark grey) branches. dN/dS values of 1, > 1, and < 1 represent neutral evolution, positive selection, and purifying selection, respectively. Arrows show the direction of selective pressure between the test and reference branches. K values < 1 or > 1 indicate relaxed or intensified selection, respectively. The Bonferroni-Holm corrected *p*-values were showed.

### Gene loss and relaxed selection on the *Vg* receptor gene (*VgR*) after *Vg* loss

Evolutionary patterns of gene loss are usually not stochastic, but rather display a clear bias with respect to gene function or genomic position^46–48^. This fact prompted us to investigate the mechanisms by which genes associated with *Vg* function evolved in *Vg*-loss lineages. We first focused on *VgR*, which is crucial for incorporation of Vg proteins into insect oocytes^38,40,41,49,50^. We hypothesized that *VgR*s may be absent or subject to distinct evolutionary pressure in *Vg*-loss species. To test this hypothesis, *VgR* genes were identified in the genomes of all 64 examined hymenopteran species. *VgR* was absent from 19 of the genomes and was present as a single-copy gene in the other 45 (Fig. 4a). Based on *VgR* presence or absence, we identified four independent *VgR* loss events throughout hymenopteran evolution; these were strongly correlated with *Vg* loss events (Fig. 4a). A high proportion of *Vg*-loss species, excluding *A. japonica*, had also lost *VgR* (78%), with only five species retaining *VgR* (namely two *Leptopilina* species, two *Trichogramma* species, and *H. didymator*). Interestingly, only one species (*Telenomus remus* of the superfamily Platygastroidea) retained *Vg* but lost *VgR*.

We next examined whether *VgR* was under relaxed selective pressure after *Vg* was lost. Using the selection intensity parameter K, where K < 1 indicates relaxed selection and K > 1 indicates strong purifying selection, we screened for signatures of relaxed selection on all test branches leading to all *Vg*-loss *VgR*-retaining species, under the RELAX model^51^ in HyPhy^52^. As expected, the result revealed a significant relaxed selection pattern on these *VgR*s (Fig. 4b; K = 0, *p* = 0). Independent tests for each of the *Vg*-loss lineages indicated significant relaxed selection on *VgR* in *Leptopilina* wasps (Fig. 4c; K = 0, *p* = 0) and in *H. didymator* (Fig. 4e; K = 0.72, *p* = 0.004), but not in *Trichogramma* spp. (Fig. 4d; K = 1.06, *p* = 0.916) or *A. japonica* (Fig. 4f; K = 1.21, *p* = 0.094). Overall, these findings suggested that *Vg* loss in endoparasitoid wasps resulted in subsequent loss or relaxed selection on *VgR*, a novel example of co-elimination of functionally linked genes^46^.

### Structural variations in the conserved VgR domain of *Leptopilina* may have led to functional loss

Relaxed selection often leads to amino acid substitutions that alter the three-dimensional structures (and therefore functions) of the encoded proteins. To determine whether this was true of VgRs in *Vg*-loss lineages, we predicted the structures of all 45 VgRs identified in this study with AlphaFold2. Based on the confidence scores (pLDDT), we focused only on the three most well-structured regions, designated A, B, and C (Fig. 5b–d). Each region contained one YWTD repeat (six YWTD motifs) and two epidermal growth factor (EGF)-like repeats (Fig. 5a). Comparative analysis suggested that the structures of these three regions were generally conserved between species (Fig. 5b–d). Notably, a portion of region A in the two *Leptopilina* VgRs differed substantially from those of all other VgRs (Fig. 5f); this region overlapped with the second EGF-like repeat of region A. Previous studies have shown that mutations in the EGF-like repeat of VgR severely disrupt the process of yolk formation^53^ and that this functional domain is essential for Vg dissociation under acidic conditions^54^. We therefore hypothesized that structural variants in this EGF-like repeat region may have led to loss of the original VgR function in *Leptopilina* species, consistent with the results of a recent study^55^.

**Fig. 5.**
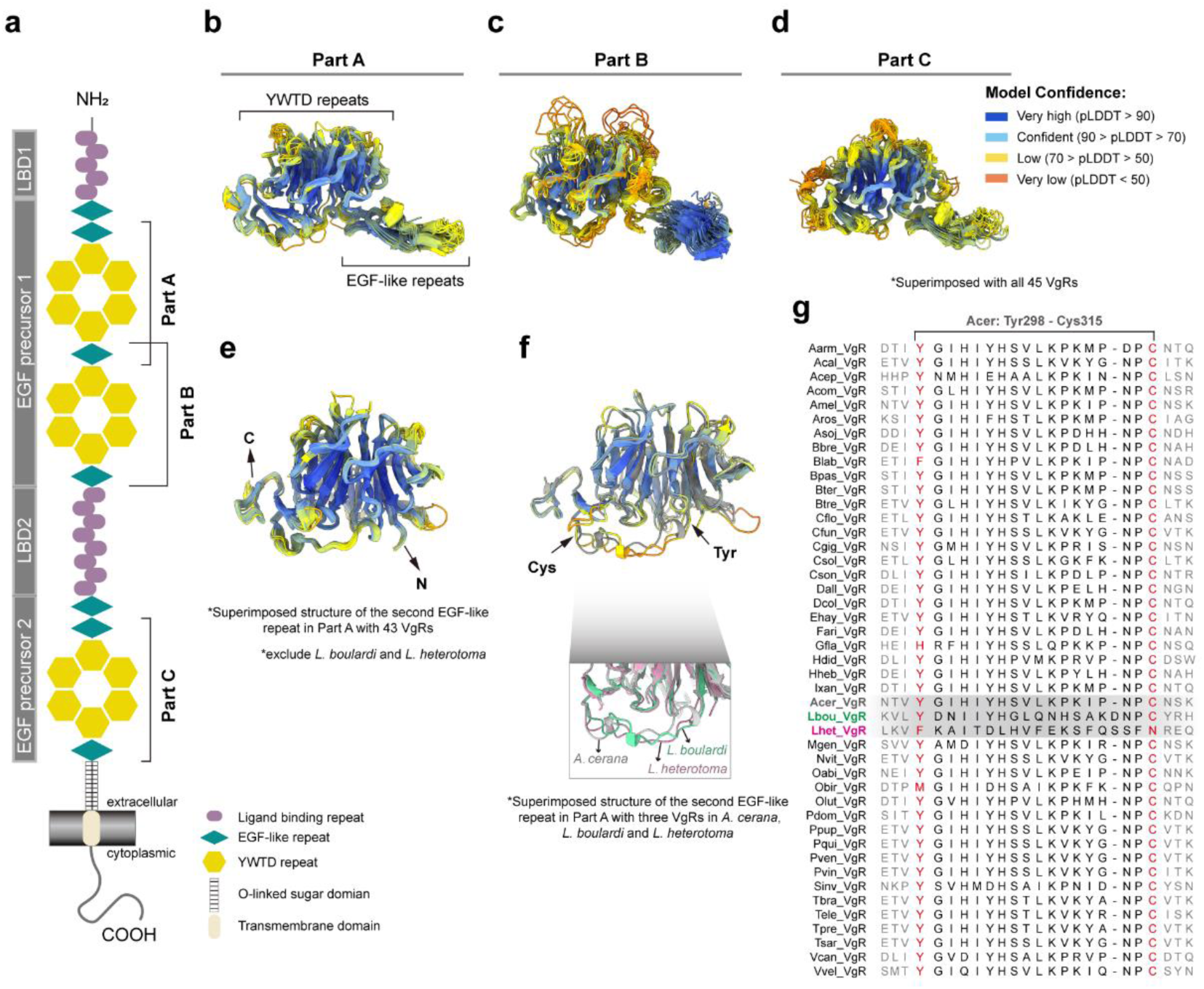
Predicted three-dimensional structures of vitellogenin receptors (VgRs). (**a**) Domain schematic of a classic VgR protein. The ligand binding domains (LBD1 and LBD2) and the EGF precursor domains (EGF precursor1 and EGF precursor2) are indicated at left. Three regions with well-defined structures based on the confidence (pLDDT) scores were designated regions A, B, and C (labeled at right). (**b–d**) Predicted structures of the VgR regions A, B, and C with AlphaFold2. Each structure is a superimposed representation based on 45 VgR sequences. pLDDT scores for each residue are indicated with color. (**e**) A superimposed structure of region A based on 43 VgRs (excluding *Leptopilina boulardi* and *L. heterotoma*) showing the highly conserved structure of the second EGF-like repeat. (**f**) Structural comparison showing variations in the second EGF-like repeat of region A in *L. boulardi* and *L. heterotoma*. (**g**) Multiple sequence alignment of the second EGF-like repeat of region A from all 45 VgRs, demonstrating the sequence variations in *L. boulardi* and *L. heterotoma*.

### The transition from ectoparasitism to endoparasitism did not directly cause *Vg*/*VgR* loss or relaxed selection

The transition from ectoparasitism to endoparasitism is considered a key step and a major trend in the evolutionary history of parasitoid wasps^23,56^. As noted above, although *Vg* and *VgR* loss events were restricted to endoparasitoids, a number of endoparasitoids retained one or both genes. This implied that the transition from ectoparasitism to endoparasitism was not the primary force driving these gene losses. To gain deeper insights into the reasons for *Vg*/*VgR* loss, we next tested whether the *Vg*s and *VgR*s in endoparasitoids evolved under relaxed purifying selection. If the transition to endoparasitism drove *Vg* or *VgR* evolution, there should be clear signals of relaxed selection on *Vg*s/*VgR*s in other *Vg-*retaining endoparasitoids compared to both ectoparasitoids and free-living species. However, the results showed no evidence of relaxed selection on *Vg*s or *VgR*s in *Vg*-retaining endoparasitoids (K = 1.04, *p* = 0.561 for *Vg*; K = 1.21, *p* = 0 for *VgR*). We therefore concluded that the ectoparasitism to endoparasitism transition was not the primary factor underlying either *Vg* loss or the subsequent loss/relaxed selection on *VgR*s in parasitoid wasps.

### Changes throughout the genome were consistent in *Vg*-loss lineages

Two genome-wide screens were next conducted to identify genetic changes in *Vg*-loss species. These screens were designed to clarify specific evolutionary patterns after *Vg* loss in different parasitoid wasp lineages and to determine whether parallel or convergent genomic changes had occurred. We first investigated changes in the sizes of gene families, also known as orthogroups (OGs). Expanded and contracted OGs were identified in the most ancient branches leading to the *Vg*-loss lineages. In each of the five independent branches associated with *Vg* loss, there were 39–462 expanded OGs and 120–713 contracted OGs (Fig. 6a).

**Fig. 6.**
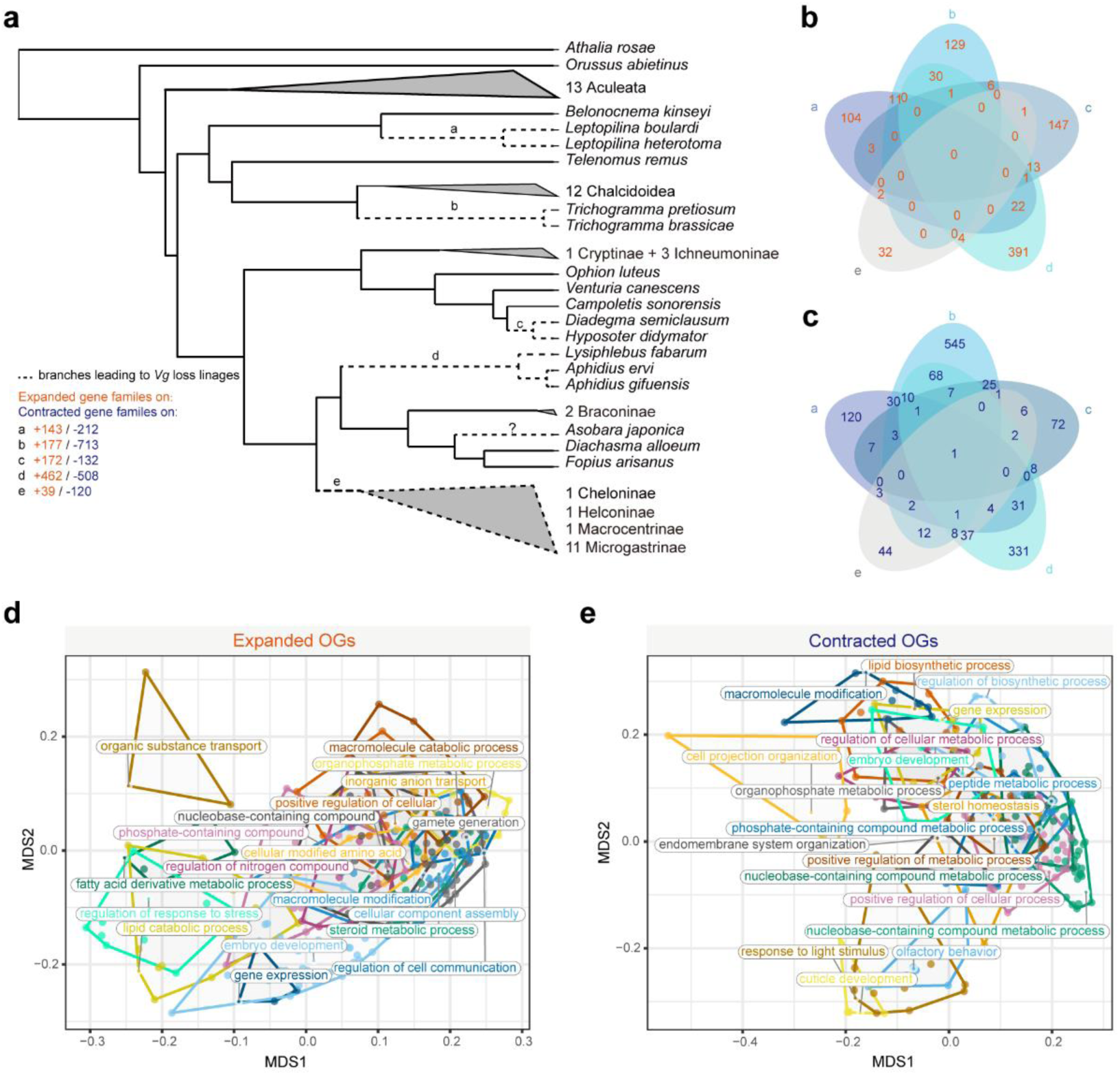
Parallel or convergent gene family evolution in the *Vg*-loss lineages. (**a**) Expansion and contraction of gene families, also known as Orthogroups (OGs) in branches leading to *Vg*-loss lineages. The phylogenetic tree was taken from Fig. 1b. The five branches affected by *Vg* loss events are indicated with dashed lines and marked a–e. (**b, c**) Unique and overlapping (**b**) expanded and (**c**) contracted OGs between the five independent *Vg*-loss branches. (**d, e**) Functional convergence of (**d**) expanded and (**e**) contracted OGs in the five independent *Vg*-loss branches (see **Methods**). Each point represents an expanded or contracted OG shared by four or five of the independent *Vg*-loss lineages. The centroid OGs were labeled with putative functions in each polygon. Gene Ontology terms associated with transport, development, and metabolism were functionally converged.

Comparative analysis showed that most changes in OG size were branch-specific; only two expanded and 40 contracted OGs were likely to have occurred in parallel (i.e., were shared between three or more lineages) (Fig. 6b, c; Supplementary Table 4, 5). The expanded OGs included a multidrug resistance-associated gene encoding a member of the ATP-binding cassette (ABC) transporter family that transports numerous substrates across the cellular membrane^57^. We inferred that parallel expansion of this OG may have enhanced the capacity of endoparasitoids to utilize host nutrients in the early development stage, compensating for the lack of nutrients from the yolk. Parallel contracted OGs included lipase, spaetzle, farnesol dehydrogenase, and trypsin.

Functional annotation was conducted for the expanded and contracted OGs between the five lineages to search for signatures of convergent functional evolution (see **Methods**). The results showed that convergently expanded or contracted OGs (those that were shared by four or all five lineages) were primarily associated with transport (e.g., organic substance transport or monoatomic ion transport), metabolism (e.g., steroid, amine, or peptide metabolic processes), and development (e.g., embryo or cuticle development) (Fig. 6d, e; Supplementary Table 6, 7).

The second genome-wide screen entailed analyzing the 2652 protein-coding one-to-one orthologous genes in the five ancestral branches leading to *Vg*-loss lineages for signatures of positive or relaxed selection (see **Methods**). One gene of unknown function was under positive selection in all five branches; five genes were under positive selection in four branches; and 27 genes were under positive selection in three branches (Supplementary Table 8). These genes were annotated with diverse functions, including “double-stranded RNA-specific editase”, “histone lysine N-methyltransferase”, and “zinc finger protein”. There were 199 genes with relaxed selection signals in all five branches.

Functional enrichment analysis indicated that these genes were related to nervous system development, embryo development, locomotion, and stress responses (false discovery rate-adjusted *p* < 0.05; Supplementary Table 9, 10; Supplementary Fig. 4). Together, these results demonstrated parallel or convergent changes in specific gene families in addition to shared gene selection events across five independent *Vg*-loss lineages. These genomic changes may have been related to adaptation to the endoparasitic lifestyle (Fig. 7).

**Fig. 7.**
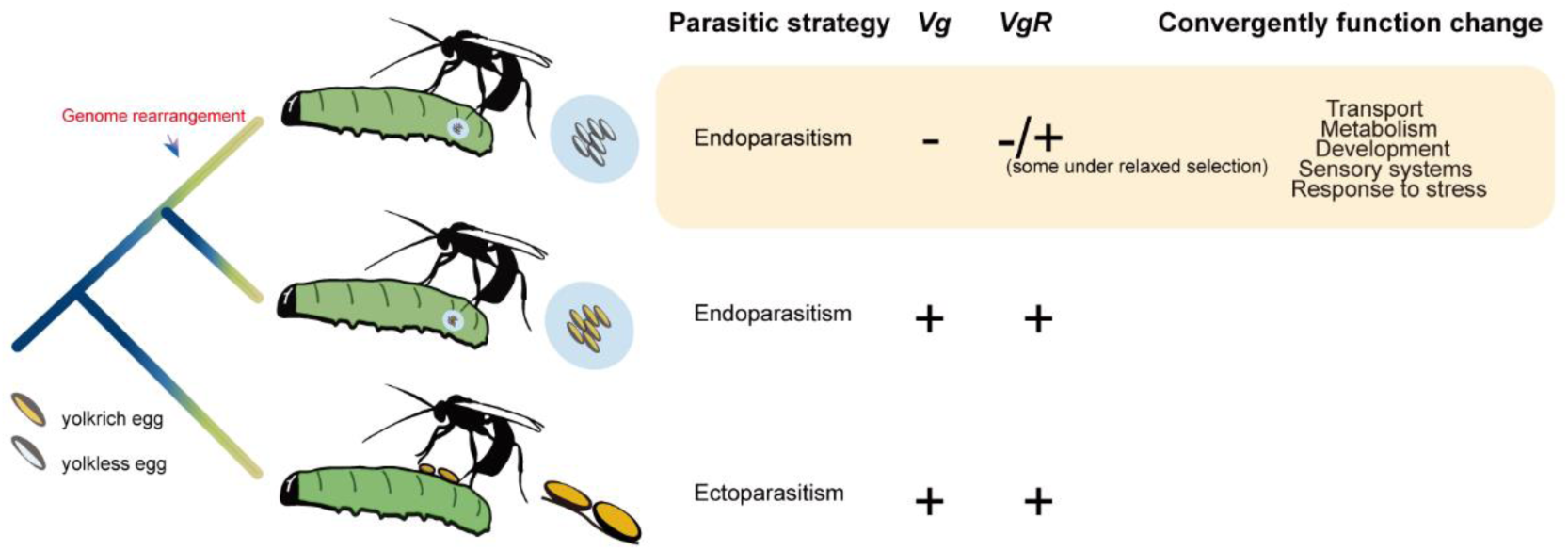
Schematic of the evolution of *Vg* and *VgR* in parasitoid wasps. The ancestral parasitoid wasp is thought to have been an ectoparasitoid that retained both the *Vg* and *VgR* genes and laid yolk-rich eggs. Hymenopteran evolution involved several independent shifts from ectoparasitism to endoparasitism. Some endoparasitoid wasps (e.g., *Pteromalus puparum*) retain *Vg* and *VgR* genes and lay yolk-rich eggs. Others have lost *Vg*, likely through genome rearrangement, leading to loss or relaxed selection of *VgR* genes due to disruption of vitellogenesis. These wasps often produce yolkless eggs (**Supplementary Table 1**). There were several shared genomic changes related to transport, metabolism, development, sensory systems, and stress resistance among the five independent *Vg*-loss lineages; these may have been related to endoparasitoid adaptation and/or contributed to loss of the yolk as an available nutrient source to the early embryo.

## Discussion

In this study, we used a comparative genomic approach to explore the genetic basis of egg yolk loss in endoparasitoid wasps, the origin and spread of which constitutes a long-standing puzzle in the field of evolutionary biology. We focused our analysis on *Vg*, an egg yolk precursor gene found in nearly all oviparous animals, including fish, birds, and insects^58^. To better understand *Vg* evolution, we annotated *Vg*s and homologs (*PVg*s and *Vgl*s) in 64 representative hymenopteran genomes. Although Vg structures and functions have been well characterized^38,41,50,59–62^, the molecular functions of PVgs and Vgls in insects are largely unknown. Here, phylogenetic and gene expression analyses revealed distinct evolutionary histories and expression patterns of *Vg* compared to *PVg*, suggesting divergent functionality of these genes. Similarly, in *Apis mellifera* and *Formica* ants, *PVg* reportedly functions in inflammation and/or oxidative stress processes rather than vitellogenesis due to sub-functionalization or neofunctionalization^63,64^. Further studies will be required in other species to clarify the functionality and evolutionary patterns of *PVg*s.

We discovered notable *Vg* loss events in 23 species from five independent lineages of endoparasitoid wasps in three superfamilies (Ichneumonoidea, Chalcidoidea, and Cynipoidea). Of the species with *Vg* loss, 14 (namely *Aphidius ervi*, *Aphidius gifuensis*, *Cotesia congregata*, *Cotesia glomerata*, *Cotesia vestalis*, *Diachasma alloeum*, *Diadegma semiclausum*, *Fopius arisanus*, *Leptopilina boulardi*, *Lysiphlebus fabarum*, *Macrocentrus cingulum*, *Microplitis demolitor*, *Microplitis mediator*, and *Venturia canescens*) have previously been identified as having yolkless eggs (Supplementary Table 1); relevant studies of egg morphology have not been conducted for the remaining 9 species. None of the species containing *Vg* have been reported to have yolkless eggs, demonstrating a strong correlation between *Vg* loss and egg yolk disappearance in the species analyzed here. In contrast to the endoparasitoids, all of the ectoparasitoids and free-living hymenopterans were found to retain at least one copy of *Vg*, demonstrating restriction of *Vg* loss to endoparasitoids.

Genome alignment and synteny analyses were conducted to identify remnants of *Vg* genes, which could provide insights into the evolutionary mechanism of *Vg* loss. Even using a low alignment threshold, *Vg* pseudogenes (e.g., those with clearly disabling insertions, deletions, frameshifts, or truncations) were not identified. This was likely due to the relatively distant timing of *Vg* loss (estimated at ∼24.32–124.58 million years ago) and the high evolutionary rates of hymenopteran genomes^65,66^. Interestingly, in three independent lineages, *Vg* loci were frequently found at genome rearrangement breakpoints, implying that a genome rearrangement event in the *Vg* region was likely the cause of gene loss in many of these species (Fig. 7). Comparison of a greater number of genomes and identification of recent *Vg* loss events would provide more robust evidence supporting this proposed mechanism of *Vg* loss.

Previous genomic studies have revealed apparent functional biases in gene loss events^46^, likely resulting from co-elimination of functionally related genes. This may be caused by changes in selective pressure on a specific trait due to adaption to a new living environment or to new biological characteristics^46,67–69^. For example, eight skin-related genes have been inactivated in both cetaceans and hippopotami over the course of evolution, potentially due to aquatic skin adaptations^70^. Fruit bats have lost functionality of three renal transporter genes (*SLC22A12*, *SLC2A9*, and *SLC22A6*) and a renal ammonium secreting transporter gene (*RHBG*), likely contributing to efficient excretion of the excess water obtained through their unique diet^71^. Similarly, we here discovered co-elimination of *Vg* and *VgR* in 21 endoparasitoid wasp species. Based on current knowledge of *Vg* and *VgR* evolution in Hymenoptera, we propose that loss or relaxed selection on *VgR* was likely a direct consequence of *Vg* loss, which rendered *VgR* dispensable (Fig. 7).

Despite the widespread loss of *VgR* among endoparasitoids, five *Vg*- loss species retained *VgR*, and the gene was under significantly relaxed selection in two *Leptopilina* wasp species (*L. boulardi* and *L. heterotoma*) and in *H. didymator*. The structure of one *VgR* functional domain was altered in the two *Leptopilina* species, suggesting a loss of the original VgR function. There are several hypotheses for *VgR* maintenance among *Vg*-loss species. First, the relatively short evolutionary time may not have been sufficient to produce inactivating mutations or complete gene loss.

Second, *VgR*s may have undergone neofunctionalization. Consistent with the latter hypothesis, a recent study of *L. boulardi VgR*, which is under relaxed selection, showed that this gene no longer performs its original function in ovary development but is involved in host-searching and mating behavior^55^. The genomic sequences of additional endoparasitoid wasps from different families and superfamilies should be analyzed in the future to further explore the coevolutionary histories of *Vg* and *VgR*.

The transition from ectoparasitism to endoparasitism may have led to relaxed selection on genes associated with nutritional status during early developmental stages (e.g., vitellogenesis genes). This is because ectoparasitoid embryos have strict nutrient requirements of the eggs (primarily the yolk proteins), whereas endoparasitoid embryos are directly exposed to a nutrient-rich external environment (the insect host hemolymph)^42,43^. Thus, the dispensability of yolk-related functions could lead to the loss of related genes in endoparasitoids. However, we found no evidence for direct selection or co-elimination of *Vg* and *VgR* as a result of the ectoparasitism to endoparasitism transition. This was supported by previous reports of functional Vg proteins in endoparasitoid wasps such as *Pteromalus puparum*^72–75^. Nevertheless, we did find that co-elimination of *Vg* and *VgR* was restricted to endoparasitoids. This may have been because endoparasitism leads to an alternative method of embryonic nutrient acquisition, which is a prerequisite for co-elimination of *Vg* and *VgR*. These two genes are crucial for formation of the yolk, which supports embryonic development in the absence of external nutrient availability. Loss of these genes could therefore lead to dependence on external nutrients, hindering a return to the ectoparasitoid state due to the generally irreversible nature of gene loss^71^.

Determination of the mechanism(s) underlying trait loss or regressive trait evolution remains challenging^46^. Here, it proved difficult to assess whether loss of the yolk protein in endoparasitoids had a neutral or positive effect on fitness. The hypothesis for positive selection suggests that yolk loss could save energy costs and increase fecundity^76^, but this hypothesis has not been directly tested. Even if loss of the yolk protein in endoparasitoids was determined to be a neutral process, other genomic changes, such as the transporter gene duplication identified through comparative genomics, could be adaptive. Detailed functional studies of closely related *Vg*-loss and *Vg*-retaining species could shed light on the impacts of *Vg* loss on fitness. In mammals, loss of the gene encoding the egg yolk protein (*VIT*) has previously been reported as coinciding with the origin of lactation and placentation^77^. We therefore propose that loss of the egg yolk is likely a consequence of regressive evolution, which may occur repeatedly during animal evolution when an alternative nutrient source becomes available during embryonic development. A study of the model species *Caenorhabditis elegans* has shown that the embryos can develop without a yolk^78^. Thus, future studies should also investigate yolk loss in nematode species, especially endoparasitic nematodes.

In conclusion, we here conducted an extensive comparative genomic analysis of 64 hymenopteran species to study loss of the vitellogenesis gene *Vg*. We found evidence of five independent *Vg* loss events throughout the order Hymenoptera, primarily occurring in species with endoparasitic strategies. *Vg* loci in species that retained the gene were frequently found in regions corresponding to genome rearrangements in *Vg*-loss species, suggesting that loss of the gene was due to genome rearrangement. Furthermore, *Vg* loss often coincided with loss or relaxed selection of *VgR*. Overall, these findings enhance our understanding of genome evolution and embryonic development across an important order of insects, revealing key genomic changes associated with yolk loss.

## Materials and methods

### Data collection

Genomic data (including genomic sequences and genome annotation results) of 64 representative hymenopteran insects were mainly collected from publicly database, including the National Center for Biotechnology Information (NCBI)^79^, InsectBase v2^80^, and Darwin Tree of Life Project^81^. For RNA-seq analysis, we also collected RNA-seq data of 10 species from NCBI Sequence Read Archive (SRA) database. Find details about data collection in Supplementary Table 11.

### *Vg* and *VgR* identification

We used Bitacora v1.4^82^ to identify candidate *Vg*s and *VgR*s. Full mode who searches genes in genomic sequences and protein annotations was selected to identify gene candidates. Besides ncbi-blast v2.13.0+ ^83^and hmmer v3.3.2^84^, the implemented tool, GEMOMA v1.7.1^85^, a homology-based gene prediction program that uses amino acid sequence and intron position conservation to reconstruct gene models from TBLASTN alignments was used to identify new genomic regions encoding gene families. We use 13 vitellogenin protein sequences of 11 species from Uniprot^86^ or NCBI database as query reference (see Supplementary Table 12). And two refence sequences of vitellogenin receptor of *Drosophila melanogaster* and *Solenopsis invicta* were retrieved from Uniprot database (see Supplementary Table 13). The specific HMM profiles were generated with alignment files of these reference sequences using hmmbuild command of hmmer v3.3.2^84^. Finally, each candidate gene was manually inspected, and further verified by checking their complete open reading frames. Then, the gene was divided into subgroups according to the following rule: sequences with N-terminal domain (Vitellogenin_N or LPD_N) as classical *Vg* (*Vg*) and partial *Vg* (*PVg*), while the other homologous members without N-terminal domain were categorized as *Vg*-like sequence (*Vgl*). For the potential chimeric sequences (e.g., potential gene model errors), we extracted the upstream and downstream 2000 bp sequences of these genes for re-annotation, using Augusts web interface^87^ (http://augustus.gobics.de/) or FGENESH+ service online^88^ (http://www.softberry.com/).

## Gene expression

We used STAR v2.7.10b^89^ and RSEM v1.3.3^90^ for transcriptomes assembly and expression calculation, respectively. Fragments Per Kilobase Million (FPKM) values were used to quantify gene expression.

## Synteny analysis

Gene synteny of selected species in three family Ichneumonidae, family Braconidae and superfamily Cynipoidea were performed with JCVI toolkit (MCScan Python version 1.3.3) ^91^ based on the alignments of each pair of species generated by comparisons from LAST^92^. And high-quality blocks of the *Vg* gene and its flanking genes were extracted for visualization using a tool implemented in the JCVI toolkit.

## Genomic rearrangement

We used LASTZ v1.04.03^93^ to perform the pairwise genome alignments with genomes of *Vg*-loss and *Vg*-retaining wasps. The *netSyntenic* files were identified with axtChain/chainNet/netSyntenic tools based on resulting alignments, and as input to identify the genomic rearrangement breakpoints followed the workflow mentioned by Liao et al^94^. We conducted a shuffling test to determine whether the median distance between the randomly shuffled breakpoints and the *Vg* locus/loci was significantly different from what would be observed. The shuffled breakpoints were produced using shuffle function of bedtools^95^ with constraint of breakpoints region on the same chromosome, and shuffling was repeated 10,000 times. The *p* value was calculated according to the method described by Sun et al.^96^.

## Gene Selection

HyPhy package v2.5.33^52^ was used for detecting gene selection signals. Indeed, aBSREL^97^ was used for detecting positively selection genes while RELAX model^51^ was used for identifying genes evolving under relaxed selection (*p* < 0.05, the Bonferroni-Holm corrected). In RELAX result, K was introduced as a parameter to assess the degree of relaxed selection, specifically K < 1 indicates that the selection was relaxed, while K > 1 indicates that the selection was strengthened. Whereas, in the result of positive selection test, the value of dN/dS greater than 1 indicated that the test branches were positively selected, otherwise they were under purifying selection.

Briefly, we used RELAX for testing relaxed selection on *Vg*s and *VgR*s. To test whether the *VgR*s of the *Vg* loss species were under relaxed selection, we first performed a general test in which all branches leading to the *Vg* loss species (excluding *A. japonica*) were set as test branches, while the remaining branches were set as reference. In addition, we also performed a separate test on each branch leading to the *Vg* loss species; To test whether the ecto-to-endo transition is a major driving force to relax the purifying selection on *Vg*s and *VgR*s and then eventually lead to their losses, we compared branches leading to endoparasitoid wasps (test) and non-endoparasitoid wasps (reference) for both *Vg*s and *VgR*s.

We also searched for common positively/relaxedly selected genes on the branches leading to the *Vg* loss lineages, which may reflect the parallel or convergent genomic changes in the *Vg* loss lineages. In total, 2652 one-to-one orthologue protein-coding genes from 17 species were involved in this selection analysis.

## Orthology inference and phylogenetic reconstruction

We utilized OrthoFinder v2.5.5^98^ to identify the orthologous and paralogous genes, the longest transcripts of genes were kept to analysis. The STAG (Species Tree Inference from All Genes) algorithm^99^ that integrated in OrthoFinder to identify Orthogroups (OGs) for phylogenetic reconstruction. Based on the usage of “-M msa” parameter, genes from 1116 OGs which were developed to leverage not only strictly single-copy orthologous genes but also the data from multi-copy gene families were used for species tree inference. Multiple sequence alignments (MSA) were inferred for each OG using MAFFT v7.520^100^ and the species tree was inferred from the concatenated MSAs using IQ-TREE v2.2.2.3^101^ with 1,000 replicates for ultrafast bootstrap analysis. The best-fitting model of sequence evolution estimated by ModelFinder that integrated in IQ-TREE v2.2.2.3^101^ was used.

And we used r8s v1.81^102^ to estimate the divergence times between species or clade. The information of divergent time was previously reported^103^ and we used two time points as fix age, 281 million years ago (mya) for Hymenoptera generated, and 8 mya for the divergent time of *Nasonia vitripennis* and *Pteromalus puparum*, and other 12 to constrain nodes marked by MCRA to calibrate the phylogenetic tree: Orussidae- Pteromalinae: 211–289 mya, Apinae-Ampulicidae: 128–182 mya, *Apis mellifera*-*Bombus terrestris*: 93–132 mya, Apinae-Microgastrinae: 203–276 mya, Apinae-Gonatopodinae: 160–224 mya, Myrmicinae-Dorylinae: 65–127 mya, Vespinae-Polistinae: 34–71 mya, Cynipinae-Telenominae: 181–246 mya, Microgastrinae-Macrocentrinae: 82–147 mya, Microgastrinae-Aphidiinae: 116–177 mya, Microgastrinae- Campopleginae: 151–218 mya, Campopleginae-Cryptinae: 79–135 mya.

## Gene family evolution

The gene family expansion and contraction were calculated using the CAFE v5^104^. The content of each OG of 64 species generated from OrthoFinder and the phylogenetic tree with divergence times inferred by r8s in above were used as input files. Based on the results, the expanded and contracted OGs of interested branches/clades that experienced *Vg* loss events were specifically extracted, and the shared OGs and associated function among these five branches/clades were inferred.

## Gene ontology enrichment and functional similarity analysis

We annotated the amino acid sequences of each species using eggNOG-mapper v.2.1.12 webserver^105^ (http://eggnog-mapper.embl.de.) against eggNOG5 database under default settings. And the biological process category of gene ontology (GO) terms was mainly focused to explore the most relevant functions in the interested OGs. In addition, GO enrichment analyses among candidate genes were performed with GOATOOLS^106^ with the go-basic.obo database (http://geneontology.org/), and p values for each GO terms were corrected for an FDR < 0.05 (Benjamini-Hochberg method of multiple testing correction). Using a simple clustering algorithm in REVIGO v1.8.1^107^, semantic relationships of enriched GO terms were analyzed and representative subset of the terms was selected for visualization.

To further unscramble the functional convergence in five *Vg* loss linages, with the aid of the R package constellatoR^108,109^ (https://github.com/MetazoaPhylogenomicsLab/constellatoR), we firstly assigned the interested OGs (expanded and contracted) onto GO terms based on the functional similarity. And the function will be clustered with an exemplar OG with the distance matrices which were transferred from pairwise OG similarity matrices.

### Protein structure analysis

We predicted the three-dimensional structures of VgR proteins from various species using LocalColabFold^110^, which implements the AlphaFold algorithm^111^. The initial multiple sequence alignments (MSAs) for each protein were generated with the MMSeqs2 server^112^. We controlled the depth of the input MSAs by setting max_msa_clusters to 512 and max_extra_msa to 1024. All five neural networks were utilized for non-template-based predictions. The max_recycle parameter was set to 3 to balance accuracy and computational efficiency, without performing the post-prediction model refinements. From the predictions generated for each species, we selected the structure with the highest internal rank for subsequent analyses. Structural alignment was performed with MatchMaker^113^ in ChimeraX^114^. A detailed list of sequences used for predictions is provided in the Supplementary Table 14. All figures were made with ChimeraX^114^.

### Data availability

All data used in this study were downloaded from public database and sources were listed in Supplementary Tables. The data supporting the findings of this study are available from the corresponding authors upon reasonable request.

## Acknowledgements

This work was supported by Key Program of National Natural Science Foundation of China (NSFC) (Grant No. 32330085 to G.Y.Y.), Program of NSFC (Grant No. 32202376 to X.H.Y.; Grant No. 32302337 to X.X.Z.), the Young Elite Scientists Sponsorship Program by China Association for Science and Technology (Grant No. 2022QNRC001 to X.H.Y.), the China Postdoctoral Science Foundation (Grant No. 2022M722804 to X.X.Z.), the Scientific Research Startup Fund Project of Zhejiang A&F University (Grant No. 2023LFR119 to X.H.Y.), the Fundamental Research Funds for Central Universities (Grant No. 2021FZZX001-31 to G.Y.Y.).

## Author contributions

X.H.Y. and G.Y.Y. conceived and designed the study. X.X.Z., Y.Y.L., C.H., Y.Y., and X.H.Y. participated in comparative genomic analysis, gene selection, genome rearrangement. K.C.C., X.X.Z., and X.H.Y. performed VgR protein structure predictions. H.W.L. and Q.F. participated in data collection and discussion. X.X.Z. and X.H.Y. wrote the manuscript with input from all of the authors. All authors read and approved the manuscript.

## Competing interests

The authors declare no competing interests.

---

**Supplementary Fig. 1.**
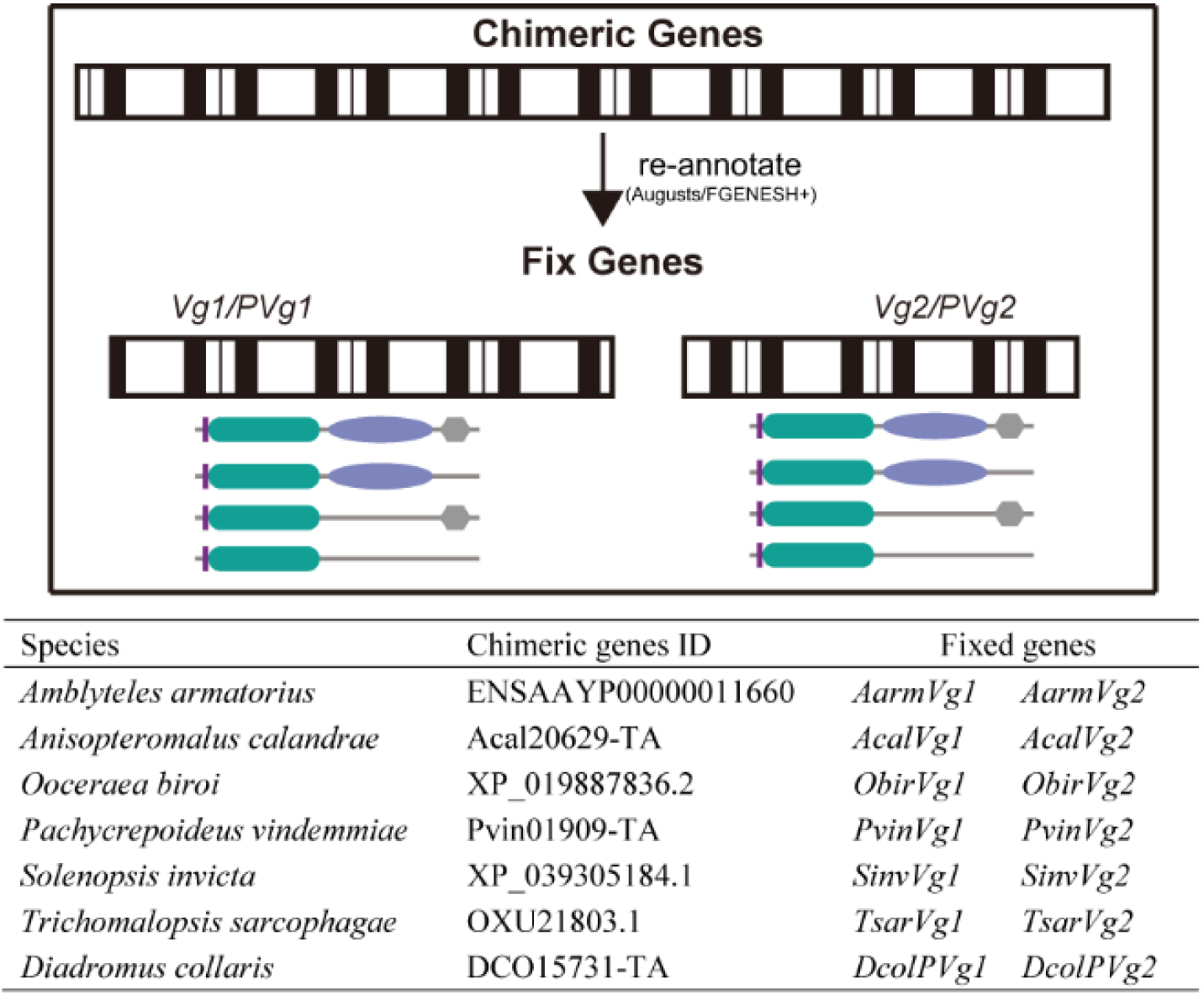
Schematic of fixation of chimeric genes. For genes joined together but owned repeat domain structures of *Vg* were split into two genes which composed domains of *Vg* or *PVg*. These genes were re-annotate using Augusts or FGENESH+ webserver, and resulted genes were further validated from using BLAST and HMMER.

**Supplementary Fig. 2.**
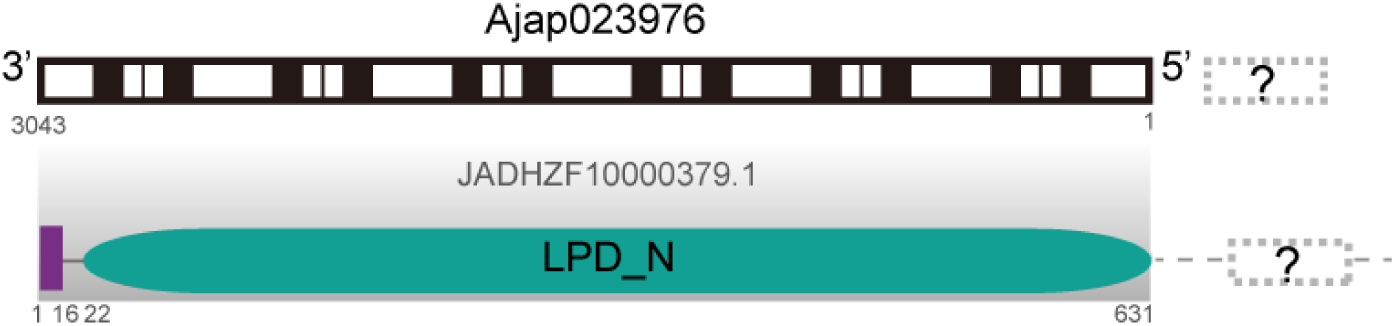
Domain structure and location of *PVg* in *Asobara japonica*. *PVg* (*Ajap023976*) of *A. japonica* was located at the start terminal on the scaffold JADHZF10000379.1 from scanning 3,043bp while the LPD_N domain predicted using SMART webserver was covered whole protein-coding sequences from 22 to 631 site with 1 to 16 was signal peptide. However, whether this *PVg* was a result of genome misassemble or bad annotation still unknown.

**Supplementary Fig. 3.**
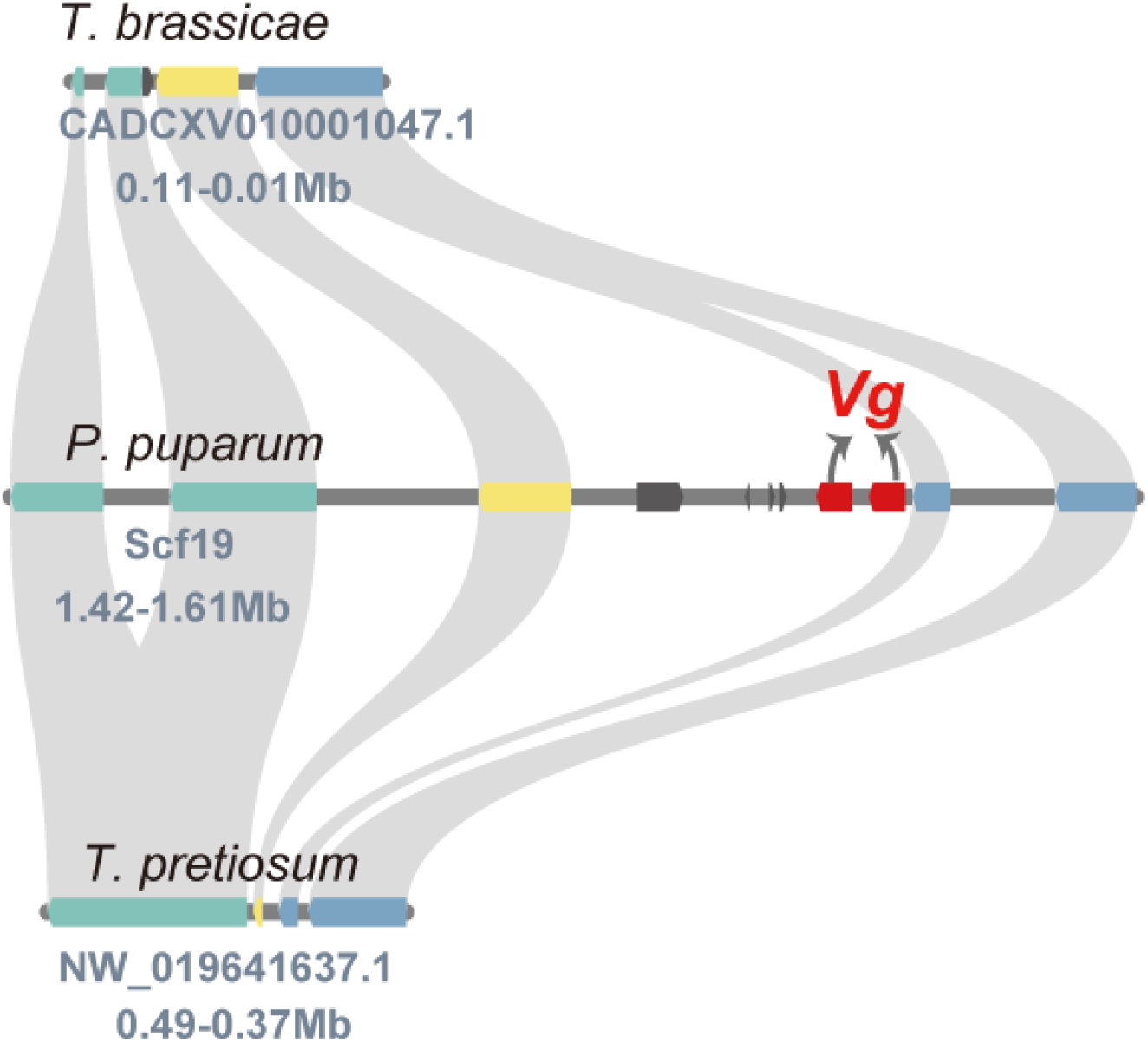
Gene synteny of *Vg*s and flanking genes among representative *Vg*-loss species and *Vg*-retaining wasps in Chalcidoidea. *Pteromalus puparum* as the *Vg*-retaining species was showed in the middle and two *Vg*s were highlighted in red, while *Trichogramma brassicae* and *T. pretiosum* were showed above and below. Results showed the two *Vg*s in *Pteromalus puparum* have no homologous genes in this syntenic block region.

**Supplementary Fig. 4.**
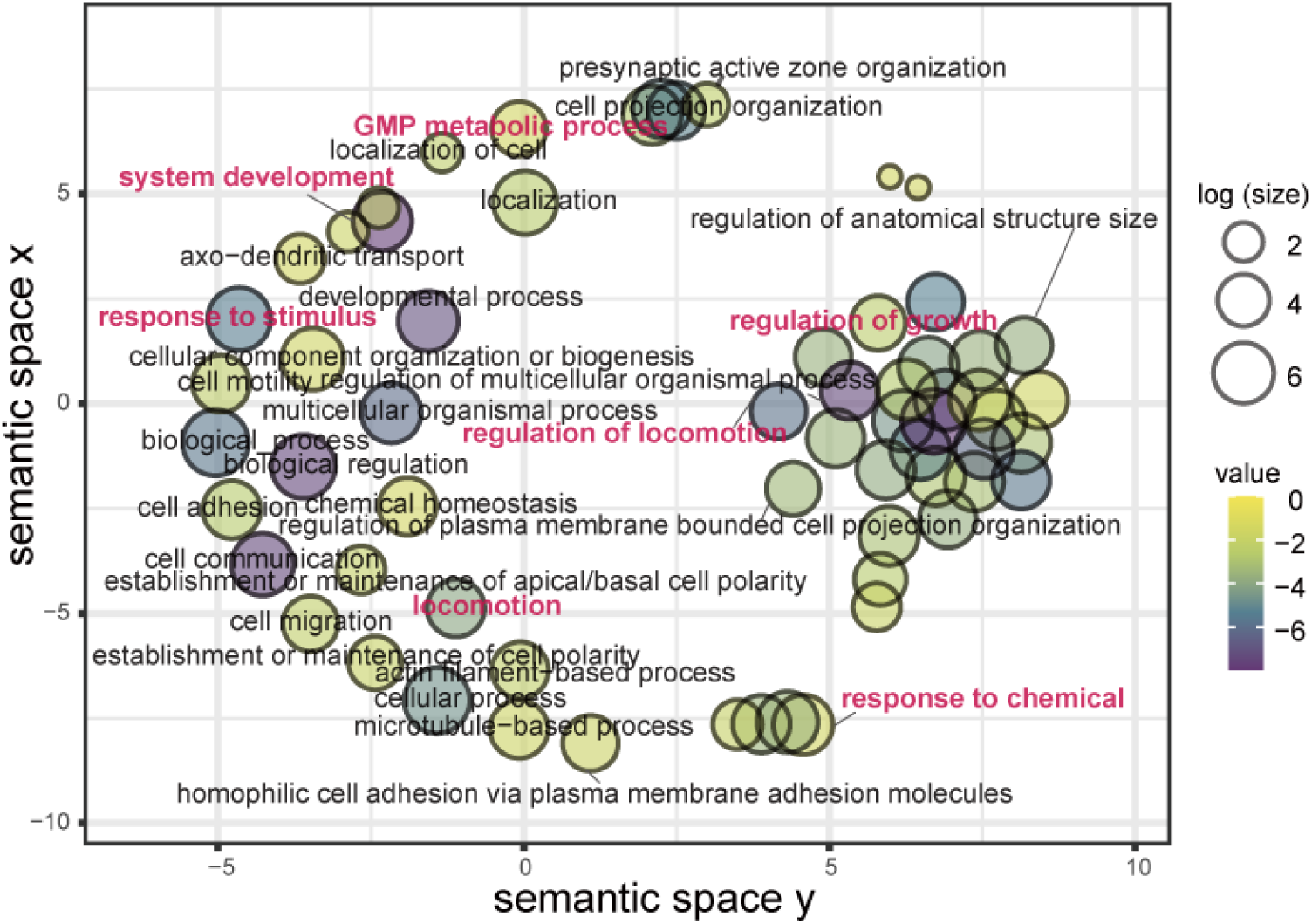
Enrichment visualization of 199 gene under relaxed selection using REVIGO. The enrichment results generated from GOATOOLS were used as input, and value C was pre-set as 0.5 which corresponding to the “small” list size, then REVIGO will clustered listed GO terms and reduce redundancy of them. Based on the information in Supplementary Table 10, dispensability of GO terms less than 0.2 were labelled in the figure and some interested GO terms were marked in pink.

## Notes

### Competing Interest Statement

The authors have declared no competing interest.

### Summary of Updates

We've polished the whole text sentence by sentence to make it more concise and standardized.

